# Non-micellar ganglioside GM1 induces an instantaneous conformational change in Aβ_42_ leading to the modulation of the peptide amyloid-fibril pathway

**DOI:** 10.1101/2023.05.12.540574

**Authors:** Manjeet Kumar, Magdalena I Ivanova, Ayyalusamy Ramamoorthy

## Abstract

Alzheimer’s disease is a progressive degenerative condition that mainly affects cognition and memory. Recently, distinct clinical and neuropathological phenotypes have been identified in AD. Studies revealed that structural variation in Aβ fibrillar aggregates correlates with distinct disease phenotypes. Moreover, environmental surroundings, including other biomolecules such as proteins and lipids, have been shown to interact and modulate Aβ aggregation. Model membranes containing ganglioside (GM1) clusters are specifically known to promote Aβ fibrillogenesis. This study unravels the modulatory effect of non-micellar GM1, a glycosphingolipid frequently released from the damaged neuronal membranes, on Aβ_42_amyloid fibril formation. Using far-UV circular dichroism experiments, we observed a spontaneous change in the peptide secondary structure from random-coil to β-turn with subsequent generation of predominantly β-sheet-rich species upon interaction with GM1. Thioflavin-T (ThT) fluorescence assays further indicated that GM1 interacts with the amyloidogenic Aβ_42_ primary nucleus leading to a possible formation of GM1-modified Aβ_42_ fibril. Statistically, no significant difference in toxicity to RA-differentiated SH-SY5Y cells was observed between Aβ_42_ fibrils and GM1-tweaked Aβ_42_ aggregates. Moreover, GM1-modified Aβ_42_ aggregates exhibited prion-like properties in catalyzing the amyloid fibril formation of both major isomers of Aβ, Aβ_40_, and Aβ_42_.

## Introduction

Pathological molecular processes involving the conformational transition of proteins/peptides into toxic oligomers and fibrils have been associated with neurodegenerative disorders such as Alzheimer’s disease (AD), Parkinson’s disease, and others[1]. In people suffering from AD, progressive accumulation of amyloid fibrils formed by the aggregation of amyloid-β (Aβ) peptides have been proposed to cause irreversible cognitive difficulties and memory loss[2]. The stochastic nature of Aβ fibrillation may lead to multiple pathways of peptide aggregation with the formation of aggregates of varying dimension (length/thickness/strength of fibrils) and secondary structure, giving rise to conformational heterogeneity and, therefore, structural polymorphism[3–8]. Increasing evidence suggests a direct correlation between these polymorphic aggregates to AD’s distinct clinical and neuropathological phenotypes [9–12]. In fact, Aβ aggregates with distinct morphologies have been observed in patients’ brains suffering from familial and idiopathic AD[13]. Moreover, pleomorphic assemblies responsible for different phenotypes observed in AD have been reinforced with the identification of differences in the morphology of Aβ fibrils generated from endogenous seeds isolated from sporadic AD and rapid-onset AD brains[14].

The environmental surroundings and other biomolecules additionally modulate Aβ aggregation. As part of the transmembrane domain of amyloid precursor protein (APP), Aβ fibrillogenesis is greatly affected by membrane lipids[15–19]. In some cases, lipid membranes provide a platform for conformational transition and subsequent generation of fibrils leading to potential disruptions of the bilayer due to increased fibril load[20,21]. Lipid-bound Aβ has also been observed among AD patients. For instance, Aβ species tightly bound to GM1 ganglioside (GM1), one of the most abundant gangliosides in the brain, was identified by Yanagisawa et al. in patients displaying early AD pathological changes[22,23]. Various in vivo and in vitro studies have reported that membrane-containing GM1 clusters act as endogenous seeds to catalyze Aβ assembly, producing toxic fibrils of varied sizes and conformations.[24–35]. Also, non-micellar GM1 lipids, mainly released from damaged neuronal membranes, affect Aβ_40_ aggregation kinetics[36,37]. Therefore, to better understand the molecular processes underlying different AD phenotypes, it is imperative to study the relationship between aggregates conformation and their pathological function, along with the mechanisms underlying strain generation and propagation in a condition closely mimicking cellular milieu.

Herein, we report that Aβ_42_, in the presence of non-micellar GM1, undergoes a spontaneous conformational transition from random coil to a predominantly soluble, β-turn and subsequently β-sheet rich intermediates before the formation of fibrillar structure. The extent and rate of structural change in Aβ_42_ alone preceding amyloid fibril formation was much lower and slow, respectively, compared to GM1-regulated Aβ_42_. Our data also indicated that GM1 interacts with the Aβ_42_ primary nucleus and forms GM1-modified fibrils. The GM1-modulated Aβ_42_ fibrils also acted as a seed for soluble, monomeric Aβ_42_ or Aβ_40_ for further formation of cross-β sheet structured fibrils.

## Materials and Methods

### Materials

GM1 (Ovine Brain, Lot: 860065P-5MG-H-021) was obtained from Avanti Polar Lipids (Alabaster, AL). Amyloid-β peptides (Aβ_40_ and Aβ_42_) were purchased from Vivitide (Gardner, MA). All other chemicals were of analytical grades and were purchased from Sigma Aldrich (St. Louis, MO) unless otherwise stated.

### Stock preparation of Aβ peptides, GM1, and ThT

GM1, Aβ_42_, Aβ_40,_ and ThT stock solutions were prepared and stored as described by Kumar et al,[38]. The critical micellar concentration (CMC) of GM1 was measured using the pyrene fluorescence assay, as reported in our recent study[36,38]. Immediately prior to use, lyophilized peptides and GM1 were dissolved in 20 mM sodium phosphate buffer, pH 7.4, to ∼200 μM and 64 μM, respectively. UV absorbance measured by nanodrop Spectrophotometer (DeNOVIX, DS-11^+^) was used to determine the exact peptide concentrations using Aβ_42_ and Aβ_40_ molecular weight as 4.5 kDa and 4.3 kDa, respectively, and 1490 M^-1^ cm^-1^ molar extinction coefficient at 280 nm. ThT concentration was determined by measuring the absorbance at 412 nm in a transparent 96-well plate (Fischer Scientific) in a Biotek Synergy 2 plate reader. The molecular weight and molar extinction coefficient of ThT dye were 318.86 Da and 36,000 M^-1^ cm^-1^, respectively.

### Real-time far-UV circular dichroism spectroscopy

The far-UV CD spectrum of 20 μM Aβ_42_ in the absence and presence of 20 μM GM1 in 20 mM phosphate buffer, pH 7.4, 37 °C was collected over a period of ∼ 22 h in a 0.01 cm path length cuvette (QS high-precision cell, Hellma Analytics) with stopper in a JASCO J-1500 CD spectrometer equipped with a Julabo AWC100 water bath following a similar method as described by Kumar et al.,[39]. The approximate scan time of each spectrum was 4.016 min, collected over a wavelength range of 180-260 nm with a scan rate of 20 nm/min, a data integration time of 2 s, a data pitch of 0.2 nm, a bandwidth of 1.5 nm, and a delay/waiting time of 0 min. The molar ellipticity of Aβ_42_ samples was smoothed using the smoothers’ function implemented within the SigmaPlot 12.0 software package.

### Determining the effect of GM1 on Aβ_42_ amyloid fibril formation via ThT fluorescence assays

Aβ_42_ (10 μM) in 20 mM sodium phosphate buffer, pH 7.4, 37 °C, was incubated in the absence and presence of varying amounts of GM1 (1 μM, 2.5 μM, 10 μM, 20 μM, and 40 μM) in 384-well microplates (Greiner bio-one black, clear bottom, non-binding, low volume, Lot number: E22053E7), with or without continuous, slow shaking (frequency = 17 Hz, i.e., 1020 cycles/min). The number of data acquisitions per hour was 10. ThT fluorescence assay was performed using below mentioned specific filters of the Biotek Synergy 2 microplate reader. A transparent sealing film was used to prevent solvent evaporation. ThT fluorescence emission was measured using a 440/40 nm filter for excitation and a 485/20 nm filter for emission. A 1:0.33 peptide:ThT molar ratio was used for all the amyloid fibril kinetic assays[40]. The fluorescence spectra of Aβ_42_ samples were plotted by subtracting the appropriate blank (that exhibited negligible ThT fluorescence) as the mean of 3 technical replicates. The blank samples had different concentrations of GM1 and ThT but no peptide. Standard error was calculated using the following equation.

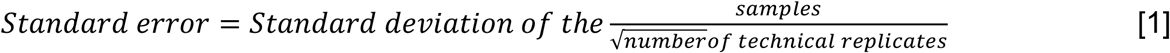

### Turbidity (optical density) measurements

The optical density at 350 nm with time was used to monitor the aggregation of 10 μM Aβ_42_ in the absence and presence of different concentrations of GM1 (1 μM, 2.5 μM, 10 μM, and 20 μM) in 20 mM phosphate buffer at pH 7.4, 37 °C under quiescent condition. The assay was conducted in 384-well microplates (Greiner bio-one black, clear bottom, non-binding, low volume, Lot number: E22053E7) sealed with transparent film. Turbidity was monitored using a Biotek Synergy 2 microplate reader. The turbidometry measurement of Aβ_42_ samples was plotted after subtracting the appropriate blank as the mean of 3 technical replicates. The standard error was calculated based on equation [1].

### Transmission electron microscopy

To obtain transmission electron microscopic (TEM) images, similar sample preparation, and TEM setting were used as described in our recent study [38]. Briefly, 5 μL sample was transferred to ultrathin holey carbon support film, copper 400 mesh grids, and negatively stained with uranyl acetate (2% w/v; Electron Microscopy Sciences) in water. TEM images were acquired using a JEOL JEM 1400 plus transmission electron microscope at a 80 kV accelerating voltage under a magnification of 25,000.

### Cell Viability Assay

Cell viability assay of Aβ_42_ amyloid fibrils was performed as described by Kumar et al.[38] Briefly, 50 μL differentiation media (Neurobasal-A; Fisher cat# 10888022, B27; Fisher cat# 17-504-044, Fisher GlutaMAX; cat# 35-050-061, Penicillin/Streptomycin) containing 60,000 SH-SY5Y cells/mL (ATCC, catalog # CRL-2266, Lot number: 63724189) were plated in a 96-well plate and differentiated for 10 days with 10 μM retinoic acid (RA) in a humidified incubator at 37 °C with 5% CO_2_[41]. Basically, differentiation media was supplemented with RA and was changed every 48 h. On the 11^th^ day, 50 μL of fresh differentiation media mixed with 50 μL of the samples containing 20 μM Aβ_42_ amyloid fibrils preformed in the absence and presence of 40 μM GM1, and 40 μM GM1 (alone) was added to the cells and incubated for 48 hours. 20 mM sodium phosphate buffer, pH 7.4, was used as the negative control, and 2% SDS was used as a positive control for cell assay. MTT cell proliferation assay (Promega, G4001) was performed to determine the toxicity of the aforementioned Aβ_42_ fibrillar samples following the manufacturer’s protocol. The average cellular viability value of the negative control (20 mM phosphate buffer, pH 7.4) after background subtraction was taken as 100%. Values reported are the average of five technical replicates, and the standard error was calculated based on equation [1] and expressed as a percentage. A Kruskal-Wallis test was performed to determine whether or not there is a statistically significant difference between the cytotoxicity of the Aβ_42_ aggregates. MTT assay was performed twice for Aβ_42_ fibrils (samples prepared from 2 different batches of Aβ_42_ peptides from the same supplier) and GM1 (same batch).

### Amyloid seeding/cross-seeding experiments

Aβ_42_ aggregates (seeds) were formed by incubating 10 μM peptide in 20 mM phosphate buffer, pH 7.4, at 37 °C, under quiescent condition, without ThT dye for 48 hours in the absence and presence of 20 μM GM1. Control Aβ_42_ samples with or without GM1, and ThT were also set up in the same microplate to track the aggregation. For seeding experiments, 5 μM of fresh Aβ_42_ in 20 mM phosphate buffer, pH 7.4, was incubated at 37 °C, under quiescent condition either without (unseeded) or with (seeded using 1% v/v, 2.5% v/v or 5% v/v) Aβ_42_ amyloid fibrils formed in the absence and presence of 20 μM GM1. For cross-seeding experiments, 5 μM of fresh Aβ_40_ in 20 mM phosphate buffer, pH 7.4, was incubated at 37 °C under no shaking condition either without (unseeded) or with (seeded using 1% v/v, 2.5% v/v or 5% v/v) Aβ_42_ amyloid fibrils formed in the absence and presence of 20 μM GM1. The aggregate formation was monitored via ThT fluorescence, as described above. Briefly, 1.7 μM (for unseeded), 1.8 μM (for 1% v/v seed), 1.95 μM (for 2.5% v/v seed), or 2.2 μM (for 5% v/v seed) of ThT was added. Fluorescence of 1.7 μM for (unseeded), 1.8 μM (1% v/v seed), 1.95 μM (2.5% v/v seed), or 2.2 μM (5% v/v seed) ThT without the fresh monomeric peptides were used as blanks, respectively. The ThT fluorescence spectra were plotted by subtracting the appropriate blank as the average value of 3 technical replicates. Standard error was calculated as mentioned above in equation [1].

## Results

### Aβ_42_ undergoes an instant conformational change upon interaction with non-micellar GM1 under Aβ_42_ amyloid forming conditions

Previous studies have shown that GM1-containing model lipid membrane induces a conformational change in Aβ upon binding to GM1 clusters[35]. We performed real-time far-UV circular dichroism (CD) of 20 μM Aβ_42_ in 20 mM phosphate buffer, pH 7.4, at 37°C with or without 40 μM of non-micellar GM1 for over 20 hours (h) (Fig. 1). As previously reported[42], fresh soluble Aβ_42_ alone exhibited a random coil structure as evidenced by the negative ellipticity peak at around 198 nm (Fig. 1A). However, after incubation of the peptide for ∼5 h under amyloid-forming conditions, a minor negative ellipticity peak at around 204 nm (indicated by the arrow in Fig. 1A) appears, suggesting random coil Aβ_42_ converting to a partially folded conformation most likely due to the formation of β-turn[43– 45]. Thereafter, the peptide transitioned to β-sheet conformation indicated by the negative peak at 218 nm (Fig. 1A, arrow). However, we noticed an immediate shift in the negative ellipticity peak of Aβ_42_ from 198 nm to ∼204 nm in the presence of non-micellar GM1 (Fig. 1B, arrow). It appears that GM1 interacts with unfolded Aβ_42_ to readily form a β-turn conformation[43–45]. After 1 h of incubation of GM1-containing Aβ_42_ in amyloid-forming conditions, the β-turn structure converted to a β-sheet rich species, as evident by the appearance of a negative minimum at 218 nm. A comparison of the magnitude and the rate of formation of β-sheet rich species in GM1-containing Aβ_42_ and Aβ_42_ alone (Fig. 1C) revealed that GM1 induces a faster and larger formation of soluble β-sheet rich species in aggregating Aβ_42_.

**Figure 1.**
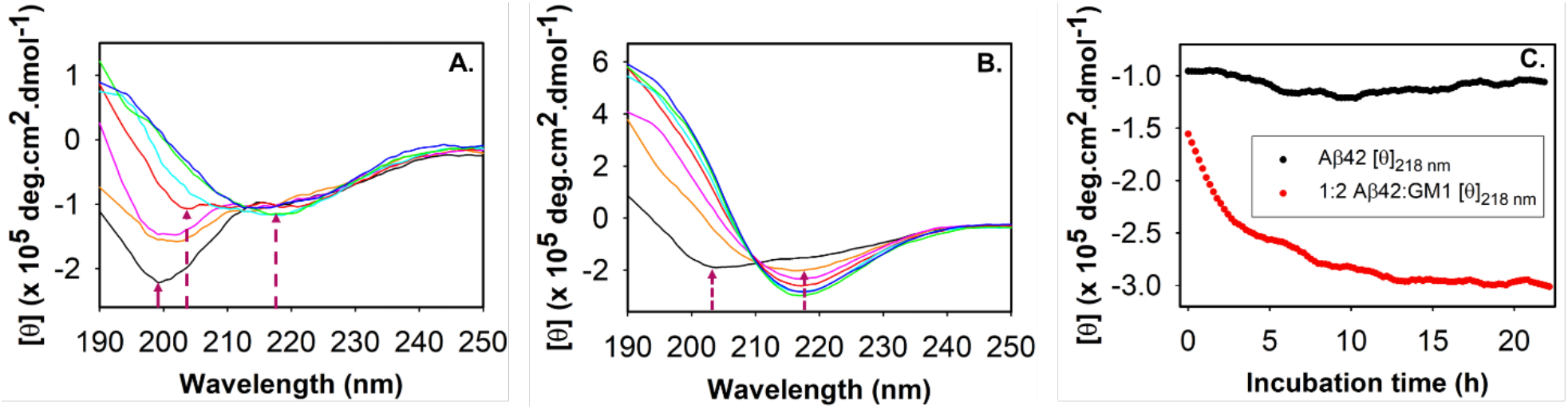
Real-time far UV-CD of Aβ_42_ with or without GM1 monitored for over 22 h. Molar ellipticity of 20 μM Aβ_42_ (A) and 20 μM Aβ_42_ with 40 μM GM1 (B) measured over a period of 22 h: black (0 h), mustard (1 h), magenta (2 h), red (5 h), cyan (10 h), green (15 h) and blue (22 h). Samples were incubated in 20 mM phosphate buffer, pH 7.4 at 37° C under quiescent condition. (C) Molar ellipticity of 20 μM Aβ_42_ (black) and 20 μM Aβ_42_ with 40 μM GM1 (red) at 218 nm over a period of ∼22 h. Arrows in (A) and (B) denote the random-coil (∼198 nm), β-turn (∼204 nm) and β-sheet (∼218 nm) peaks.

### GM1 interaction with the Aβ_42_ amyloidogenic nucleus likely produce GM1-modified fibrils

To understand the change in Aβ_42_ amyloid fibril formation propensity due to the GM1-induced conformational change in the peptide, we performed a ThT fluorescence assay on 10 μM Aβ_42_ alone and in the presence of different concentrations of GM1 (Fig. 2A) in 20 mM phosphate buffer, pH 7.4 at 37 °C, under constant shaking. Aβ_42_ spontaneously formed amyloid fibrils resulting in an exponential polymerization curve with the absence of a distinct lag phase suggesting that under shaking, the peptide undergoes a nucleation-independent mechanism of protein aggregation. In the isodesmic or linear polymerization mechanism, fibrillation proceeds via a sequence of multiple steps, where the successive addition of amyloidogenic monomer to the growing fibril is energetically favorable without the need of a nucleus[39,46,47]. Increasing concentrations of GM1 up to a 4-fold molar excess to Aβ_42_ did not affect the rate of Aβ_42_ fibril formation. However, a minimal diminution in ThT fluorescence was observed. Transmission electron microscopy (TEM) images of 10 μM Aβ_42_ alone and in the presence of 2.5 μM GM1 acquired after 24 h incubation in the above-mentioned conditions confirmed the presence of amyloid fibrils all over the TEM grid (Fig. 2B, Fig. 2C). Interestingly, when the fibrillation of 10 μM Aβ_42_ was slowed down by incubating it at pH 7.4, 37 °C, and without shaking with the introduction of a lag phase, a significant decrease in ThT fluorescence was observed with increasing GM1 concentration (Fig. 2D). The lag phase represents a period where the primary nucleus is generally formed from monomers [47]. Both 20 μM and 40 μM GM1 showed a similar and maximum decline in ThT binding profiles. Therefore, we used a 1:2 molar ratio of Aβ_42_ :GM1 throughout the study. Moreover, the larger reduction in ThT binding, when Aβ_42_ aggregated slowly in a typical nucleation-dependent polymerization compared to the spontaneous nucleation-independent process, likely suggests that non-micellar GM1 interacts with the primary Aβ_42_ nucleus in addition to the monomer to inhibit amyloid fibril formation. To confirm this, we obtained TEM images of 10 μM Aβ_42_ with or without 20 μM GM1 incubated for 48 h in physiologically relevant pH, temperature, and quiescent condition (Fig. 2E, Fig. 2F). On the contrary to the reduced ThT binding; we observed fibrillar species of Aβ_42_ formed in the presence of GM1 all over the TEM grid similar to that of Aβ_42_ alone. The Aβ_42_ sample incubated with GM1 appeared to have shorter fibrillar species along with the presence of oligomers compared to the control Aβ_42_ (only) sample. To further investigate the contrasting findings from ThT fluorescence assays and TEM images, we measured the optical density at 350 nm of Aβ_42_ with or without GM1 (Fig. 2F). A trivial increase in the turbidometry measurements of the peptide samples with increasing concentration of GM1 indicated that the extent of Aβ_42_ aggregates formed in the absence and presence of GM1 is somewhat similar. Moreover, no drastic change in the Aβ_42_ lag phase was observed with or without GM1 (Fig. 2D), suggesting that the glycosphingolipid does not prevent the formation of the peptide amyloidogenic nucleus. These data suggest that GM1 lipids most likely interact with the primary nucleus of Aβ42 and regulate it to form shorter fibrillar species that probably bind less efficiently to ThT.

**Figure 2.**
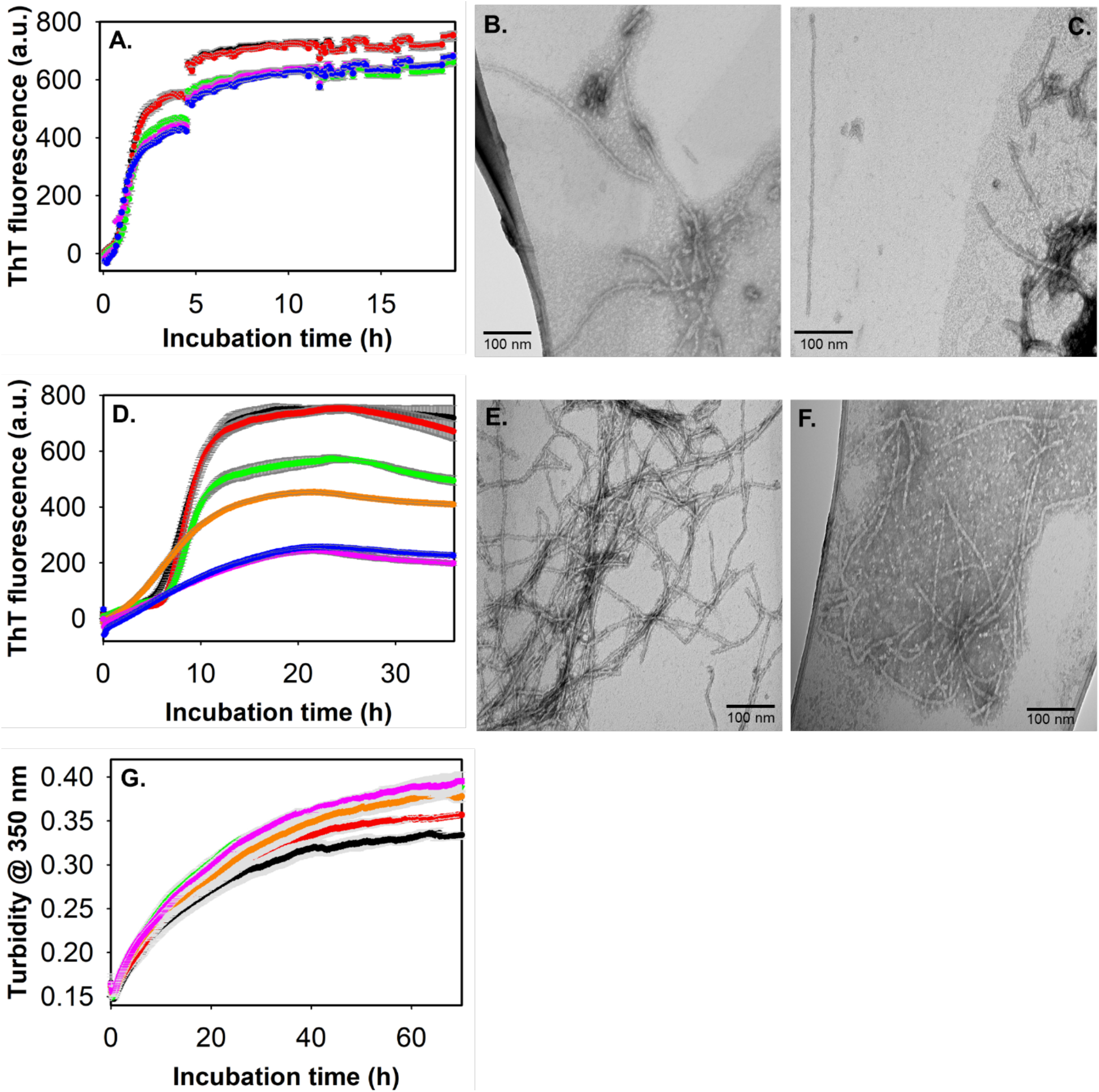
Effect of GM1 on Aβ_42_ amyloid fibril formation. (A) ThT fluorescence intensity of 10 μM Aβ_42_ (black), and with varying concentrations: 1 μM (red), 2.5 μM (green), 20 μM (magenta), and 40 μM (blue) of GM1 under slow continuous shaking condition in 20 mM phosphate buffer, pH 7.4 and 37° C. TEM images of 10 μM Aβ_42_ alone (B) and in the presence of 2.5 μM GM1 (C) acquired after 24 h incubation at physiological pH, temperature, and constant shaking conditions. (D) ThT fluorescence intensity of 10 μM Aβ_42_ (black), and with varying concentrations: 1 μM (red), 2.5 μM (green), 10 μM (mustard), 20 μM (magenta), and 40 μM (blue) of GM1 under quiescent condition in 20 mM phosphate buffer, pH 7.4 and 37° C. ThT fluorescence intensity in (D) was scaled to match the highest ThT fluorescence intensity in (A) by multiplying by the factor of 1.6. TEM images of 10 μM Aβ_42_ alone (E) and in the presence of 20 μM GM1 (F) acquired after 48 h incubation at physiological pH, temperature, and no shaking condition. Scale bar as indicated in the images. (G) Optical density measurement of 10 μM Aβ_42_ (black) and in the presence of GM1: 1 μM (red), 2.5 μM (green), 10 μM (mustard), and 20 μM (magenta) at 350 nm in 20 mM phosphate buffer, pH 7.4 at 37° C, no shaking as a function of time. Data are presented as the average ± standard error, calculated from 3 replicates (n = 3). Standard error bars are represented in grey in (A), (D), and (G).

### Statistically, no significant difference was observed in the cytotoxicity of GM1-modified Aβ_42_ fibrils and Aβ_42_ fibrils to human neuroblastoma cells

The process involving the conversion of the soluble non-toxic Aβ peptides to toxic form is believed to be a critical step in the development and progression of AD, especially when toxic effects of GM1-containing membranes mimicking lipid rafts catalyzed Aβ fibrils, are well documented[27,48]. So, we evaluated the cytotoxic effect of non-micellar GM1-modified Aβ_42_ fibrils on human neuronal SH-SY5Y cells. The cell viability assay on differentiated human neuroblastoma SH-SY5Y cells treated with GM1-modulated Aβ_42_ fibril appeared to be slightly toxic (∼18%) (Fig. 3). Samples containing Aβ_42_ fibril and GM1 did not show any significant cytotoxicity on RA-differentiated SH-SY5Y cells (Fig. 3). The p-value, calculated by Kruskal-Wallis test is 0.35. Hence the cytotoxicity of Aβ_42_ aggregates formed in the absence and presence of GM1 was found to be insignificant at p < 0.05.

**Figure 3.**
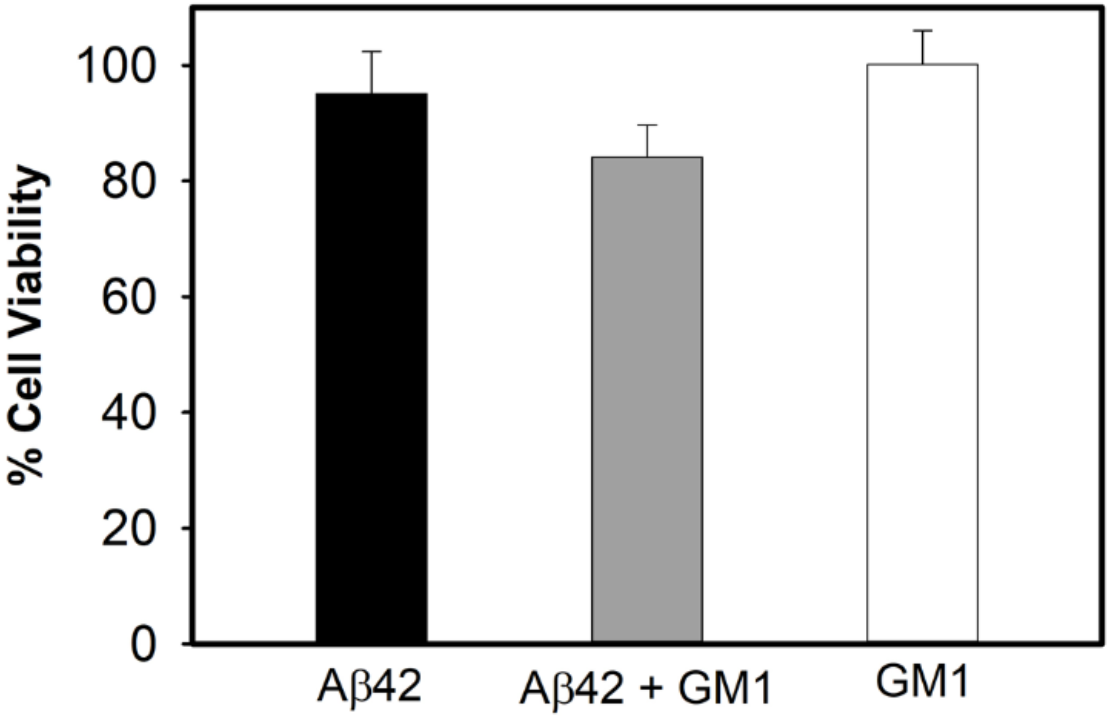
Evaluation of the neurotoxic effect of Aβ_42_ amyloid fibrils. The MTT assay was performed with RA-differentiated human neuroblastoma SH-SY5Y cells to determine their metabolic activity as an indicator of cell viability upon treatment with solutions containing Aβ_42_ aggregates formed from 10 μM Aβ_42_ without (black) and with 20 μM GM1 (grey). Measurements were carried out after 2 days of incubation. The white bar is of 2-day old 20 μM GM1 alone. Data are presented as the average ± standard error, calculated from 5 replicates (n = 5). Kruskal-Wallis test indicates that the result is insignificant at p < 0.05.

### GM1-modified Aβ_42_ fibrils can catalyze the aggregation of soluble monomeric Aβ_42_ as well as Aβ_40_

Besides the direct cytotoxic effect exhibited by the aggregates, the deleterious fibril surfaces can catalyze self and neighboring peptides/proteins to aggregates via secondary nucleation[49]. Self-seeded (homogeneous) and cross-seeded (heterogeneous) secondary nucleation is monomer-dependent processes whereby transient monomer binding from the same peptide/or neighboring protein, respectively, to a fibril lateral surface accelerates aggregation leading to fibril proliferation and fibril load[50]. We, therefore, tested the seeding and cross-seeding effects of un-sonicated Aβ_42_ fibrils using ThT fluorescence assays (Fig. 4). The secondary nucleation processes of Aβ_42_ were studied by adding different amounts of preformed Aβ_42_ fibrils to 5 μM of soluble unfolded Aβ_42_ monomers under quiescent condition at physiological pH and temperature (Fig. 4A). The addition of 1%, 2.5% or 5% v/v preformed Aβ_42_ fibrils as seed led to an immediate increase in ThT fluorescence, suggesting spontaneous Aβ_42_ fibril formation. Meanwhile, the addition of 1%, 2.5%, or 5% v/v preformed Aβ_42_ fibrils to 5 μM of soluble unfolded Aβ_40_ monomers significantly reduced the lag-phase of 5 μM Aβ_40_ aggregation kinetics (Fig.4B). Seeding and cross-seeding of Aβ_42_ and Aβ_40_, respectively, have earlier been observed with sonicated Aβ_42_ fibrils[51]. An immediate rise in ThT fluorescence was also observed when GM1-modified Aβ_42_ fibrils were used for seeding. However, the use of GM1-free Aβ_42_ fibrils as seed resulted in faster rate kinetics (Fig. 4C). On the other hand, catalysis of 5 μM Aβ_40_ aggregation using GM1-modified Aβ_42_ fibrils as seed (for cross-seeding) required a higher seed amount (5% v/v) (Fig. 4D). Other noticeable difference observed in Aβ_42_ aggregates’ seeding/cross-seeding activities is significant less ThT fluorescence intensity in the case of GM1-modified Aβ_42_ fibrils compared to GM1-free Aβ_42_ fibrils. Secondary nucleation in GM1-modified Aβ_42_ fibrils appears to be less efficient in catalyzing the Aβ peptides. This may be due to GM1 bound to Aβ_42_ fibrils, or perhaps GM1-modulated Aβ_42_ fibrils, produced Aβ_42_ and Aβ_40_ aggregates with altered conformations which bind less efficiently to ThT.

**Figure 4.**
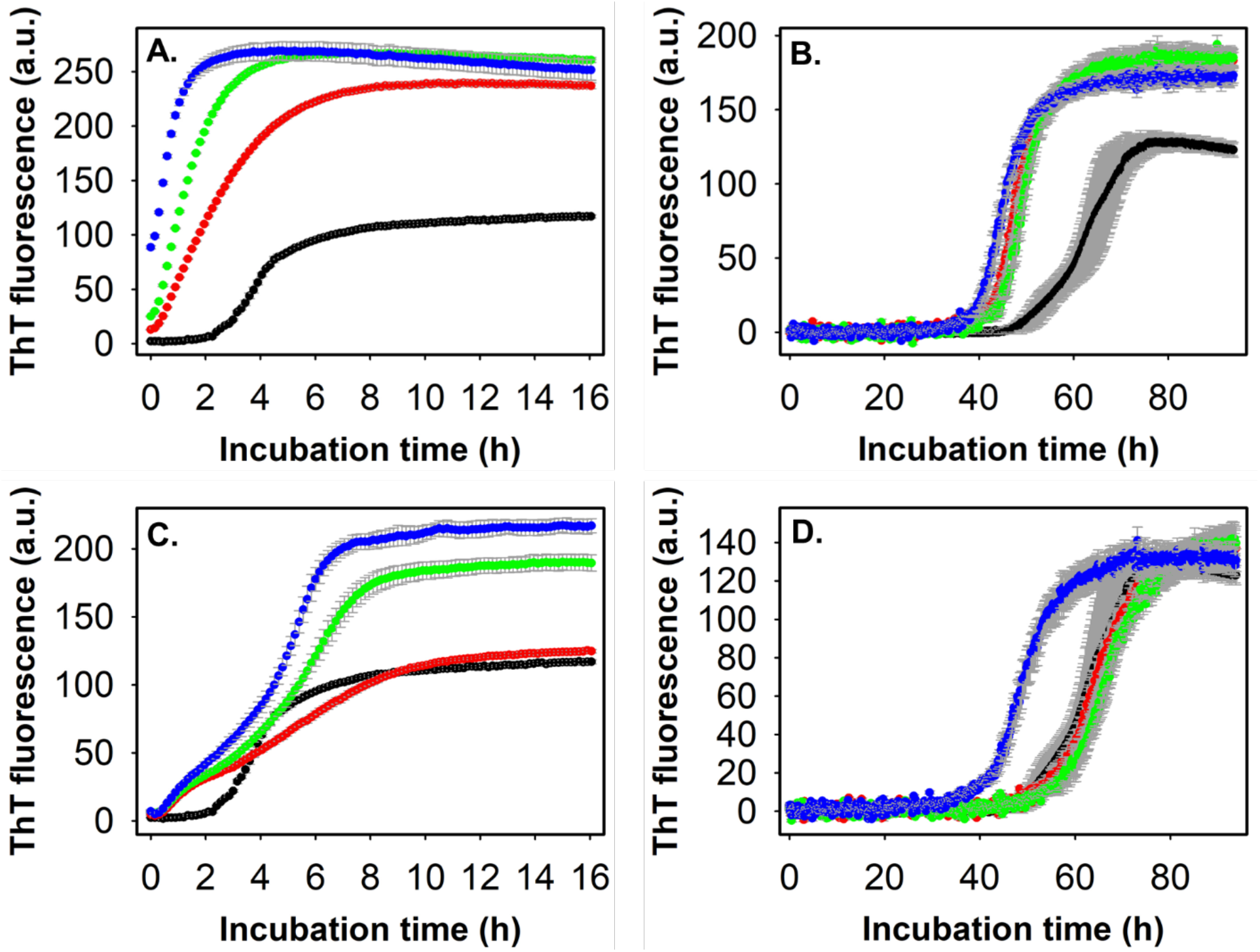
Seeding and cross-seeding of Aβ_42_ and Aβ_40_ with pre-formed Aβ_42_ aggregates. (A) ThT fluorescence profiles of 5 μM Aβ_42_ alone (black) and in the presence of 1% v/v (red), 2.5% v/v (green) and 5% v/v (blue) pre-formed Aβ_42_ amyloid fibrils in 20 mM phosphate buffer, pH 7.4, 37° C under quiescent condition. (B) ThT fluorescence profiles of 5 μM Aβ_40_ alone (black) and in the presence of 1% v/v (red), 2.5% v/v (green) and 5% v/v (blue) pre-formed Aβ_42_ amyloid fibrils in 20 mM phosphate buffer, pH 7.4, 37° C under quiescent condition. (C) ThT fluorescence profiles of 5 μM Aβ_42_ alone (black) and in the presence of 1% v/v (red), 2.5% v/v (green) and 5% v/v (blue) pre-formed GM1-modified Aβ_42_ fibrils in aforementioned conditions. (D) ThT fluorescence profiles of 5 μM Aβ_40_ alone (black) and in the presence of 1% v/v (red), 2.5% v/v (green) and 5% v/v (blue) pre-formed GM1-modified Aβ_42_ fibrils in above mentioned conditions. An appropriate blank was subtracted in each case. Data are presented as the average ± standard error, calculated from 3 replicates (n = 3). Standard error bars are represented in grey.

## Discussion

The deleterious self-association of intrinsically disordered monomeric Aβ forms large fibrillar aggregates that deposit as insoluble protease-resistant plaques in AD. NMR-based studies revealed structural differences between fibrils propagated in vitro using brain-derived amyloid seeds as templates from different AD patients[11,52]. Similarly, variations in Aβ fibril structure in vivo were identified using conformation-sensitive fluorescent dyes and antibodies in the brain that correlate with diverse phenotypes in AD[53,54]. In analogy to prion strains associated with distinct clinical and pathological phenotypes, different Aβ fibril conformation is believed to be responsible for familial or sporadic rapidly progressing AD[9,55–57]. Aβ peptides are known to form polymorphic fibrils with variations in molecular structure and toxicity depending on fibril growth conditions and interaction with other biomolecules such as lipids, including GM1-containing model lipid membranes[58–62].

GM1, the most abundant brain ganglioside, is clustered in plasma membranes region known as lipid rafts, but free GM1 is frequently released from damaged neuronal membranes[63,64]. Herein, we systematically investigated the effects of non-micellar GM1 on the amyloid aggregation of Aβ_42_. As schematically summarized in Figure 5, at physiological pH, temperature, and quiescent condition, non-micellar GM1 lipids interact with intrinsically disordered Aβ_42_ monomers and spontaneously form a dominant β-turn fold before transforming to β-sheet rich species and subsequently forming GM1-modified fibrils. Similar structural conversions of Aβ_42_ in hexafluoroisopropanol (HFIP) have previously been reported[43]. To better understand the role of GM1-induced structural transformations in Aβ_42_, we monitored the aggregation propensity of the peptide in the presence of GM1. We noticed that GM1 significantly reduced ThT fluorescence intensity when Aβ_42_ aggregates in a quiescent condition at physiological pH and temperature.

**Figure 5.**
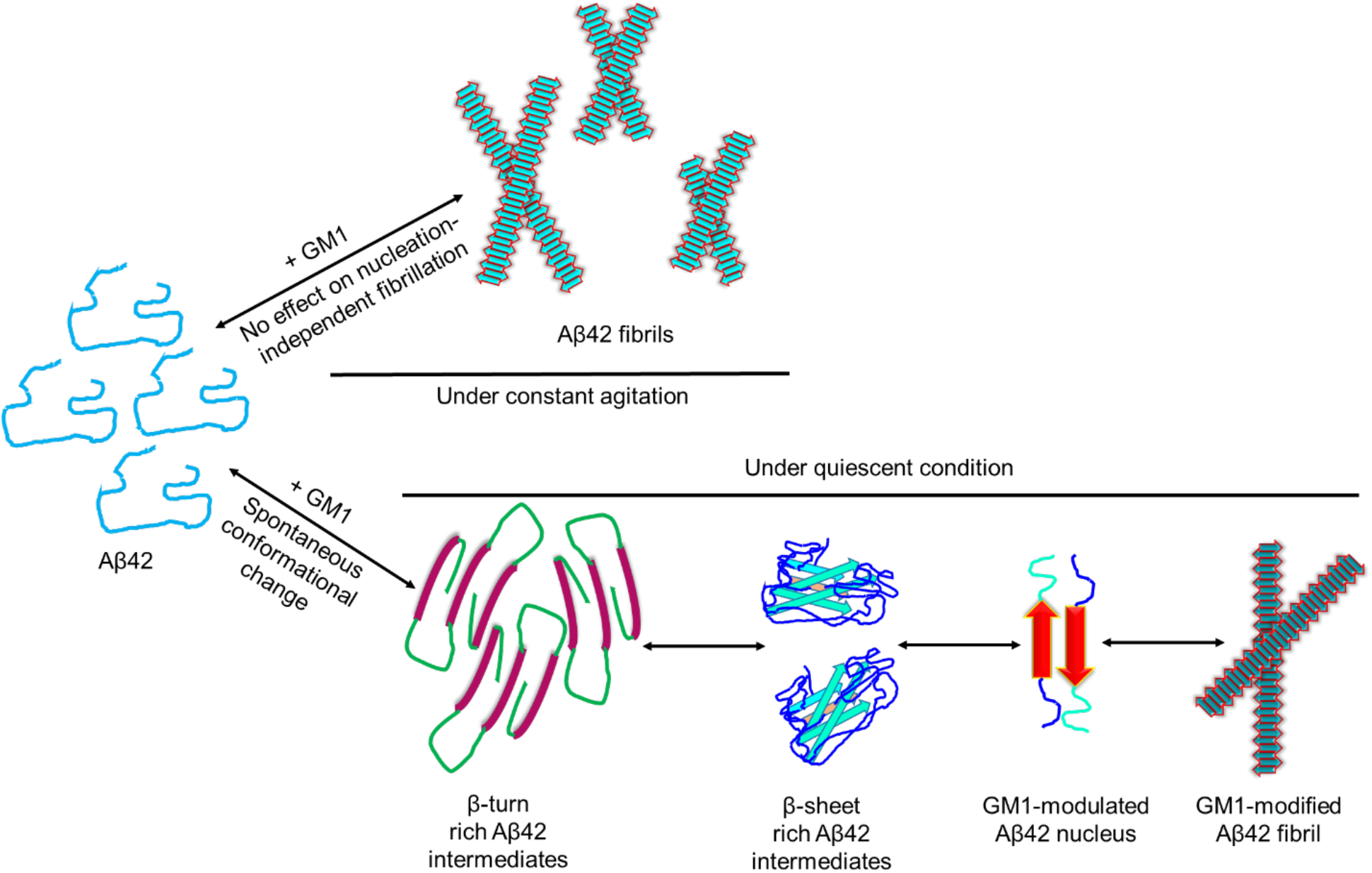
Schematic representation of the effect of non-micellar GM1 on Aβ_42_ fibrillation. At physiological pH, temperature, and quiescent condition, non-micellar GM1 spontaneously converts random-coil Aβ_42_ to a dominant β-turn and then to β-sheet rich species. Upon subsequent generation of the Aβ_42_ amyloidogenic primary nucleus, GM1 additionally interacts with amyloidogenic nucleus to produce GM1-modified Aβ42 nucleus, and ultimately GM1-modulated fibrils. Non-micellar GM1 fails to regulate Aβ42 amyloid fibril formation when the peptide instantaneously converts to amyloid fibril via nucleation-independent polymerization under constant agitation at physiological pH and temperature.

However, a minimal effect was observed under constant agitation when the peptide forms amyloid fibril instantaneously. Under non-shaking conditions, amyloid fibril formation of Aβ_42_ is a nucleation-dependent polymerization, where the soluble monomers transition to form an amyloidogenic precursor, often known as the primary nucleus, which subsequently forms amyloid fibrils. Under the constant shaking condition, the conversion of monomeric Aβ_42_ to amyloid fibrils is instantaneous and follows nucleation-independent polymerization without the need of a primary nucleus. These observations suggest that in the absence of the Aβ_42_ primary nucleus, GM1 is ineffective in modulating the peptide fibrillation. This observation is in agreement with a recent biophysical study by Chakravorty et al.[36]. Slowly aggregating Aβ_42_ under quiescent condition generates the primary nucleus and provides sufficient time for non-micellar GM1 to interact with the primary nucleus and to affect the fibril formation[47,65].

The ability of non-micellar GM1 to rapidly convert random-coil Aβ_42_ monomers to β-turn, and β-sheet rich intermediate species, and interact with the peptide primary nucleus perhaps led to the formation of GM1-modified Aβ_42_ fibrils with an altered structure as indicated by less ThT binding (∼4-fold) compared to Aβ_42_ fibrils formed in the absence of GM1. The enhanced fluorescence exhibited by ThT upon binding to a typical amyloid fibril is mainly attributed to the presence of cross-β sheet architecture in the fibril[66–70]. A substantially reduced ThT fluorescence intensity in GM1-modulated Aβ_42_ fibril suggests inefficient binding of the fluorescent amyloid marker, possibly due to altered cross-β architecture. Fibril conformational differences have been identified before using ThT binding characteristics[71,72]. The other explanation for the reduced ThT fluorescence intensity of Aβ_42_ in the presence of GM1 compared to Aβ_42_ alone, may be due to the presence of a significant amount of soluble oligomers and fewer amyloid fibril species. Indeed, far-UV CD data suggests greater extent of β-sheet rich Aβ42 species formation in the presence of GM1 than in the absence of GM1. Nonetheless, non-micellar GM1 lipids interact with various species of Aβ_42_ and modulate the peptide amyloid fibril pathway leading to the formation of GM1-modified fibrils. To further characterize the biochemical properties of GM1-modulated Aβ_42_ fibrils, we tested their toxicity (direct) and secondary nucleation abilities (in-direct toxicity). GM1-modified Aβ_42_ fibrils, and Aβ_42_ fibrils formed in the absence of GM1 statistically exhibited no significant difference in direct toxicity towards RA-differentiated human neuroblastoma cells (SH-SY5Y). Additionally, secondary nucleation is a process that self-catalyzes (seeding) or accelerates neighboring peptides/proteins (cross-seeding) to aggregate formation leading to fibril proliferation. Our results show that the GM1-modified Aβ_42_ fibrils could accelerate monomeric Aβ_42_ (via seeding) and monomeric Aβ_40_ (via cross-seeding) aggregation. However, the aggregates bound less efficiently to ThT, possibly due to altered cross-β architecture. The spread of pathogenic inclusions in many neurodegenerative diseases, including AD, akin to a ‘prion infectious nature’ can be attributed to the accelerated conversion of non-aggregated proteins/peptides through primary and secondary nucleation [50,73,74].

The results presented here demonstrate that Aβ amyloid pathway can be modulated upon interaction with other biomolecules, including lipids in the surrounding cellular milieu. Furthermore, discoveries of GM1-modified Aβ fibrils can unequivocally establish the differences in Aβ aggregates that may lead to different AD phenotypes. From a therapeutic perspective, preventing the formation of biomolecules regulated Aβ aggregates resembling those described here could be a valuable strategy in combating neurodegenerative disorders, including AD.

## Abbreviations

(AD): Alzheimer’s Disease
(Aβ): Amyloid-β
(GM): Ganglioside
(ThT): Thioflavin T
(CD): Circular Dichroism
(TEM): Transmission Electron Microscopy
(MTT): 3-(4,5-dimethylthiazol-2-yl)-2,5-diphenyl-2H-tetrazolium bromide)

## Acknowledgment

This study was supported by the National Institutes of Health Grants (AG048934 and DK13221401 to A.R.).

## Author’s Contribution

M.K. and A.R. designed the research; M.K. performed the experiments and processed the data; M.K., M.I. and A.R. analyzed the data; M.K., M.I. and A.R. wrote the manuscript; and A.R. directed the project.

